# Lipid Droplets are active translation depots that respond to infection

**DOI:** 10.1101/2025.05.12.653387

**Authors:** Rakesh Mohan Jha, Dilip Menon, Arpita Nahak, Akhil Kumar, Vineet Choudhary, Sheetal Gandotra

**Affiliations:** CSIR-Institute of Genomics and Integrative Biology, Sukhdev Vihar, New Delhi-110025, India; Academy of Scientific and Innovative Research (AcSIR), Ghaziabad-201002, India; Department of Biotechnology, All India Institute of Medical Sciences, Ansari Nagar, New Delhi-110029, India

## Abstract

Lipid droplets (LDs) are increasingly recognized as dynamic organelles with roles extending beyond lipid storage. While the core LD proteome across cells reveals minimal overlap, several proteins with “housekeeping” functions such as ribosomal proteins and translation factors have largely been ignored as contaminants. Here, we provide biochemical and microscopy-based evidence of ribosome components and the ribosome to be present in association with the LD. By developing a modified method based on puromycylation, we demonstrate that active ribosomes are localised in close-proximity of LDs, demonstrating that LDs serve as sites of active protein translation. By using *Mycobacterium tuberculosis* (Mtb) infection as a model, we show that LD-associated translation is regulated under infection conditions. Although the specific mRNAs translated on LDs remain to be identified, these findings reveal LDs as critical platforms for protein synthesis and provide insights into how the LD proteome is established. This work highlights the multifaceted role of LDs in cellular homeostasis and host-pathogen interactions.

**IMPORTANT:**

- Manuscripts submitted to Review Commons are peer reviewed in a journal-agnostic way.
- Upon transfer of the peer reviewed preprint to a journal, the referee reports wil be available in ful to the handling editor.
- The identity of the referees wil NOT be communicated to the authors unless the reviewers choose to sign their report.
- The identity of the referee wil be confidentia ly disclosed to any a filiate journals to which the manuscript is transferred.

**GUIDELINES:**

- For reviewers: https://www.reviewcommons.org/reviewers
- For authors: https://www.reviewcommons.org/authors

**CONTACT:** The Review Commons o fice can be contacted directly at

## Introduction

Lipid droplets (LDs) are specialized subcellular organelles that originate at the endoplasmic reticulum and become distinct entities that can traffic to other regions of the cell and continue to make independent contacts with organelles such as mitochondria, peroxisomes, and lysosomes, in addition to the ER (Jacquier *et al*, 2011; Kien *et al*, 2022; Benador *et al*, 2018; Chang *et al*, 2019; Menon *et al*, 2023; Schulze *et al*, 2020; Song *et al*, 2022; Freyre *et al*, 2019). The neutral lipid core is surrounded by a phospholipid monolayer, is further decorated by a variety of proteins (Thiam *et al*, 2013). While some of these proteins regulate lipid metabolism, some are involved in organelle-organelle contacts, trafficking, gene regulation, and some are thought to be simply stored there (Mejhert *et al*, 2020; Li *et al*, 2012; Bersuker *et al*, 2018). Stimuli such as infection, innate immune activation, and hormonal stimulation lead to differential recruitment of proteins to the surface of LDs, in a cell type dependent manner (Rösch *et al*, 2016; Menon *et al*, 2019; Bosch *et al*, 2020; Brasaemle *et al*, 2004; Anand *et al*, 2012). Despite differences in LD proteomes across conditions and cell types, some proteins have repeatedly been found yet ignored as “contaminants” during organelle isolation; ribosomal proteins belong to this class of proteins.

Ribosomes are the most abundant sub-cellular complexes, with a large density associated with the cytosolic face of the ER membrane. More recently, localized translation on peroxisomes, on mitochondria, and within the nucleus has been documented (David *et al*, 2012; Dahan *et al*, 2022; Zipor *et al*, 2009; Lesnik *et al*, 2015; Williams *et al*, 2014). Localized translation has several advantages; (i) economical transport of a single mRNA compared to thousands of proteins, (ii) provision of a local high density of the protein that is destined for its function at that site, rather than relying on diffusion-dependent or specialized trafficking-dependent mechanisms for achieving requisite concentrations of the protein, (iii) regulated translation at specific sites based on stimulus and (iv) avoiding mis-localization of proteins (Bourke *et al*, 2023; Unsworth *et al*, 2010; Lashkevich & Dmitriev, 2021).

Currently there are two principal pathways for targeting proteins to lipid droplets. One is the cytosol-to-LD, or the CYTOLD pathway, wherein proteins are synthesized in the cytoplasm and then bind directly bind to LDs. The other is ER-to-LD, or the ERTOLD pathway, wherein these proteins are initially inserted into ER membrane and later transported to LDs (Olarte *et al*, 2022). While the CYTOLD pathway relies on amphipathic helices, lipid anchors and protein-protein interactions to associate with LDs, the ERTOLD pathway relies on migration of the membrane inserted helical regions of monotypic assembly to the LD monolayer membrane. These models are insufficient to explain LD localization of several proteins, such as ribosomal proteins, suggestive of indirect binding. In addition, the increase of ribosomal protein density in the LD proteome upon infection, suggests that ribosomal proteins may have a concerted action to play at the LD during infection (Menon *et al*, 2019). Whether these ribosomal proteins are part of an active ribosomal complex, performing moonlighting functions or are simply stored on lipid droplets is not understood.

In this article we use a combination of biochemical and microscopy-based approaches to demonstrate that ribosomal components as well as active ribosomes are present on LDs. We also report that these translation events are modulated upon infection by the intracellular pathogen *Mycobacterium tuberculosis*. This discovery introduces a distinct mode of targeting of newly synthesized proteins to LDs, by localized translation at the LDs. These findings expand our current knowledge of the function of LDs, to platforms that are actively engaged in protein synthesis.

## Results

### Ribosomal components associate with lipid droplets

Ribosomes are primarily composed of ribosomal proteins, rRNAs and other proteins known as ribosomal associated proteins. Several studies have detected ribosomal proteins and translation factors in LD proteome (Sato *et al*, 2006; Wan *et al*, 2007; Zhang *et al*, 2011; Yu *et al*, 2015; Rösch *et al*, 2016; Menon *et al*, 2019; Mejhert *et al*, 2020). A comprehensive list of these studies has been summarized in Supplementary table 1. LDs are closely associated with the endoplasmic reticulum (ER), facilitating the exchange of proteins and lipids across these membranes. Due to the high ribosomal content of the ER membrane, there is a substantial risk of ER-derived contamination during LD isolation (Song *et al*, 2022; Jacquier *et al*, 2011). To assess the presence of ribosomal components, and ensure the purity of isolated LDs, protein markers were screened using immunolocalization on LDs isolated from either COS7 cells or THP1 macrophages (Figure 1A). This approach revealed presence of PLIN3, an LD coat protein, but the lack of the ER-marker Calreticulin from the surface of the LD (Figure 1B). To detect the ribosomal components, RPL4-a component of its large subunit, was chosen as it is present at the peptide exit center and surface accessible in the ribosomal complex (Ban *et al*, 2000; Wilson *et al*, 2020). In addition, RPS9-a component of the small subunit, was chosen as it was previously identified by multiple studies in the LD proteome (Menon *et al*, 2019; Mejhert *et al*, 2020). Both the antibodies labeled discrete regions on the surface of LDs, confirming their association with LDs (Figure 1C). While no background of RPL4 was seen in these preps, RPS9 positive puncta separated from the LDs were also visible, underscoring the need for a microscopy-based approach for validating proteins found in the LD fraction.

**Figure 1:**
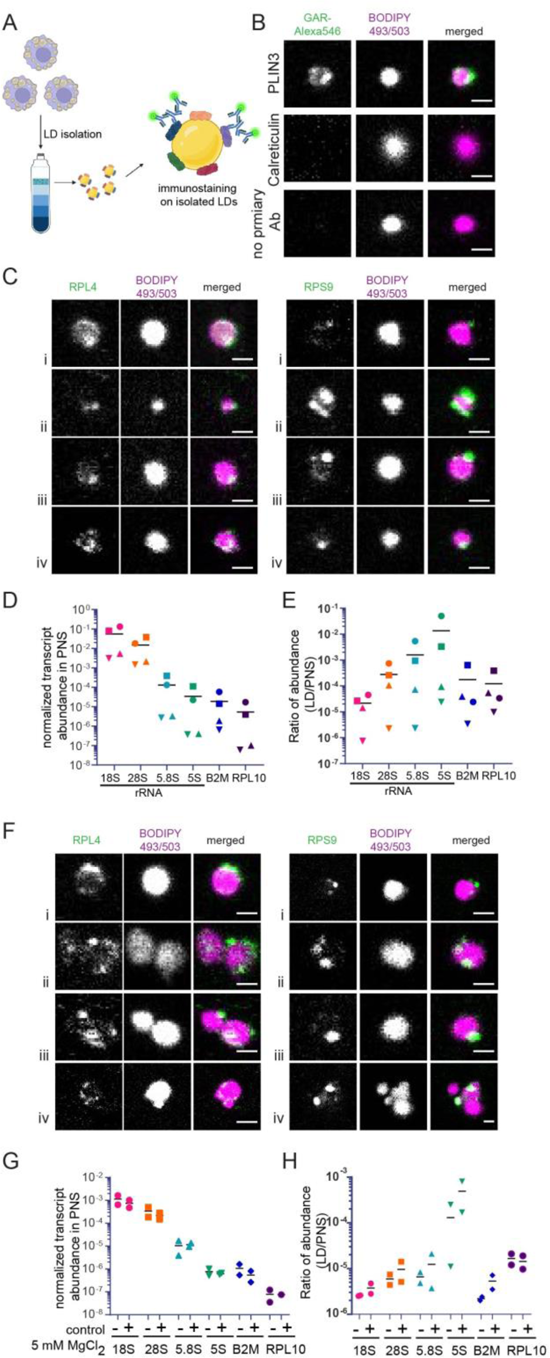
Presence of rRNAs and ribosomal proteins on lipid droplets: (A) Schematic representation of the lipid droplet (LD) enrichment and immunostaining process. (B) Confocal images of LDs isolated from COS7 cells and labelled with BODIPY 493/503 and immunostained for PLIN3, and calreticulin. (C) Multiple representative confocal images of LDs isolated from THP1 macrophages and labelled with BODIPY493/503 and immunostained for RPL4 and RPS9. Scale bar: 1 μm. Data are representative of two independent experiments. (D, E) Transcript abundance of rRNAs, B2M, and RPL10 genes in the post-nuclear supernatant (PNS) (D) and lipid droplet (LD) fraction, normalized to PNS (E), measured by qRT-PCR. N = 4 independent experiments. (F) Confocal images of lipid droplets labelled with BODIPY 493/503 and immunostained for RPL4 and RPS9 after purification using 5 mM MgCl₂-containing buffer. Scale bar: 1 μm. (G, H) Transcript abundance of rRNAs, B2M, and RPL10 genes in the PNS and LD fraction measured by qRT-PCR after lipid droplet isolation using a high MgCl₂-containing buffer (N =2).

To test for the presence of rRNA, qRT-PCR was performed after RNA isolation from the isolated LD fraction. We chose multiple housekeeping genes, B2M, RPL10, UBE2D2 and EEF1A, to select for those with total cellular abundance similar to rRNA genes. The abundance of B2M and RPL10 was found to be similar to 5.8S and 5S rRNAs in the total RNA and chosen for inclusion in these experiments (Figure 1D). We reproducibly found the presence of all rRNAs as well as some mRNAs in the LD fraction (Figure 1E). The relative abundance of 5S and 5.8S rRNA was on average 322.2±268.5 times and 40.9±27.2 times higher than that of 18S rRNA. Even 28S rRNA, a component of the large subunit was 9.2±2.8 times more abundant than 18S rRNA. This was despite higher abundance of the 18S rRNA, a component of the small ribosomal subunit by several orders of magnitude higher than the large subunit rRNA (Figure 1D). Interestingly, relative enrichment of the gene RPL10 was also similar to that of 28S and 18S rRNA, despite lower abundance in the total cellular RNA (Figure 1D and 1E). Due to the low abundance of RNA in the LD fraction, we initially used 15 T175 flasks for LD isolation. However, subsequent studies with only a single T175 flask also enabled detection of rRNA in the LD fraction, despite 100-folds lower levels of rRNA in the post-nuclear supernatant (Figure 1F-G). Importantly, even in this experimental strategy, relative enrichment of 5S rRNA over other transcripts was observed (Figure 1G). As RNA structure and binding to other biomolecules is dependent on divalent metal ions, we questioned if addition of 5 mM MgCl2 to the LD isolation buffers would increase our ability to detect rRNA. Addition of MgCl2 neither altered detection, nor enrichment of these transcripts in the LD fraction (Figure 1G and H). Furthermore, this change did not alter detection of RPS9 and RPL4 on the LD surface (Figure 1H). This raised the possibility of inherently tight association between LDs and ribosomes, dependent on protein binding, rather than direct binding of rRNA.

RPs and rRNAs could be present on LDs in a translation-dependent or independent manner. We reasoned that disruption of the translation complex should result in loss of detection of ribosomal proteins and rRNA on LDs if they are present in a translation-dependent manner. Divalent cations are crucial for ribosome stability, and it has been demonstrated that high concentrations of EDTA (15 mM) can disrupt polysomes (Nousch *et al*, 2014; Ron *et al*, 1968). So, we used a buffer containing high concentrations of EDTA and EGTA for LD isolation. Despite isolation of LDs with such a buffer, RPS9 and RPL4 were still detected associated with LDs (Figure 2A), suggesting even monosomes to be associated with LDs. Additional, non-LD associated RPS9 puncta were readily detectable under this condition (Figure 2Aii and iv). Puromycin is a global translation inhibitor that is expected to dissociate the active translation complex but not disrupt the individual ribosomal subunits (Azzam & Algranati, 1973). Retention of RPS9 and RPL4 on LDs despite puromycin treatment suggested the association of ribosomal proteins/subunits in a translation-independent manner, in addition to the possibility of a translation dependent manner (Figure 2B).

**Figure 2:**
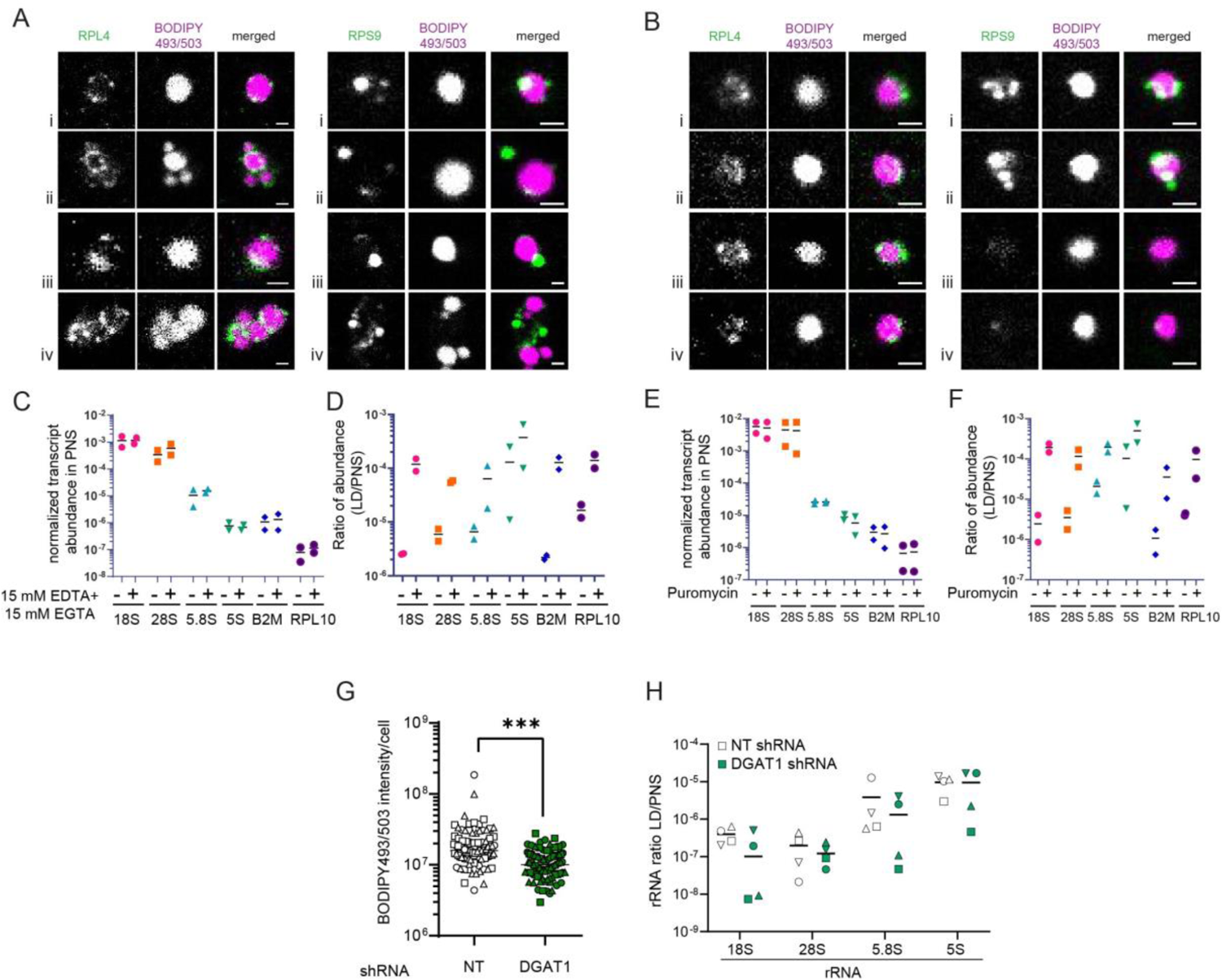
Ribosomal proteins on LDs and rRNAs in the LD fraction can also be independent of translation and lipid droplets (A, B) Multiple representative confocal images of lipid droplets labelled with BODIPY 493/503 and immunostained for RPL4 and RPS9 after purification with high EDTA/EGTA concentration buffers (A) or puromycin-treated cells (B). Scale bar= 1 μm. Four representative images of individual LDs are shown. Data are representative of two independent experiments. (C-F) Transcript abundance of rRNAs, B2M, and RPL10 genes in the post-nuclear supernatant (PNS) (C and E) and lipid droplet (LD) fraction, normalized to PNS (D and F) measured by qRT-PCR (N=2). Panels C and D are from experiments where LDs were isolated with high EDTA and EGTA (15 mM each) buffers and Panels E and F are from experiments where LDs were isolated after puromycin treatment of macrophages. (G) Signal quantification for LipidTox DeepRed-stained THP1 macrophages stably expressing non-targeting (NT) or DGAT1-targeting shRNA. Data are on a per-cell basis, pooled from 3 independent experiments where a minimum of 25 cells were imaged per experiment. Symbols from the same experiment are represented by the same shapes. (****p < 0.0001, from a two-tailed Mann-Whitney test). Scale bar: 10 µm. (H). Transcript abundance of all rRNAs in LD fraction of THP1 cells expressing non-targeting (NT) or DGAT-1 targeting shRNA, measured by qRT-PCR. Symbols from the same experiment are represented by the same shapes (N = 4).

Interestingly, the abundance of rRNAs increased in the LD fraction upon treatment with either 15 mM EDTA/EGTA (Figure 2D) or Puromycin (Figure 2F) without impacting total levels of these RNAs (Figure 2C and 2E). Interestingly, both B2M and RPL10 levels in the LD fraction also increased by an order of magnitude with both these treatments. This suggested that conditions that favor dissociation of ribosomal subunits or polysomes, enable easier detection of RNA in the LD fraction. This could be due to these transcripts or their protein-bound complexes simply floating in the LD fraction. We argued that this could be tested by quantification of rRNA in the top fraction from cells expressing DGAT1 shRNA or control shRNA. As reported previously (Jaisinghani *et al*, 2018), DGAT1 silencing significantly reduced LD abundance in THP1 macrophages (Figure 2G). In 3 out of 4 independent experiments, we observed 2-10 folds lower abundance of 18S and 5.8S (2 experiments for 5S rRNA) in the top fraction from DGAT1 silenced macrophages (Figure 2H). However, there was no significant difference in rRNA in the LD fraction upon DGAT1 silencing, suggesting that while ribosomal proteins do selectively localize to the LD, rRNA or a nucleoprotein complex containing rRNA could also independently float to the top fraction during LD isolation. Therefore a fractionation based approach on its own is insufficient to conclusively state that ribosomes associate with LDs.

### Lipid droplets host active ribosomes

We performed transmission electron microscopy of THP1 macrophages to visualize ribosomes. Negatively stained sections revealed that ribosomes associated with some but not all LDs. In some instances, while the ribosomal density was visible on the surface of LDs without an associated ER membrane (Figure 3A), ER-associated ribosomes were also found to be associated with LDs in other cases (Figure 3B). To test whether these ribosomes are translationally active, we used the ribopuromycylation (RPM) method (David *et al*, 2012). This method involves the use of two translational inhibitors-puromycin and emetine. Puromycin terminates translation by covalent attachment to the C-terminus of the elongating peptide while emetine is an elongation inhibitor that freezes ribosomes in the elongation state. As a result this combination locks the newly synthesized puromycylated peptide in the ribosomes (David *et al*, 2012). We used an antibody against puromycin to detect the sites of active translation in the cells. However, since translation occurs throughout the cells, the RPM signal was detected throughout the cell (Supplementary Figure 2). To assess LD-localized puromycylation, we combined RPM with proximity ligation assay (Hegazy *et al*, 2020) (RPM-PLA) and checked the proximity of ribosomes containing puromycelated peptides with PLIN2, the major LD coat protein in THP1 macrophages (Figure 3C). As a positive control for this assay, we used antibodies of mouse and rabbit origin against the PLIN2. In this case multiple high intensity PLA puncta were present around LDs (Figure 3D, middle panel). The combination of anti-puromycin (rabbit) and anti-PLIN2 (mouse) antibody generated fewer number of PLA puncta but these were indeed in the proximity of LDs (Figure 3D, top panel). Omission of one of these antibodies led to the complete abrogation of the PLA reactivity (Figure 3D, bottom panel). These data reveal that RPM-PLA can be used as a reliable readout for LD-proximal ribosomal activity.

**Figure 3:**
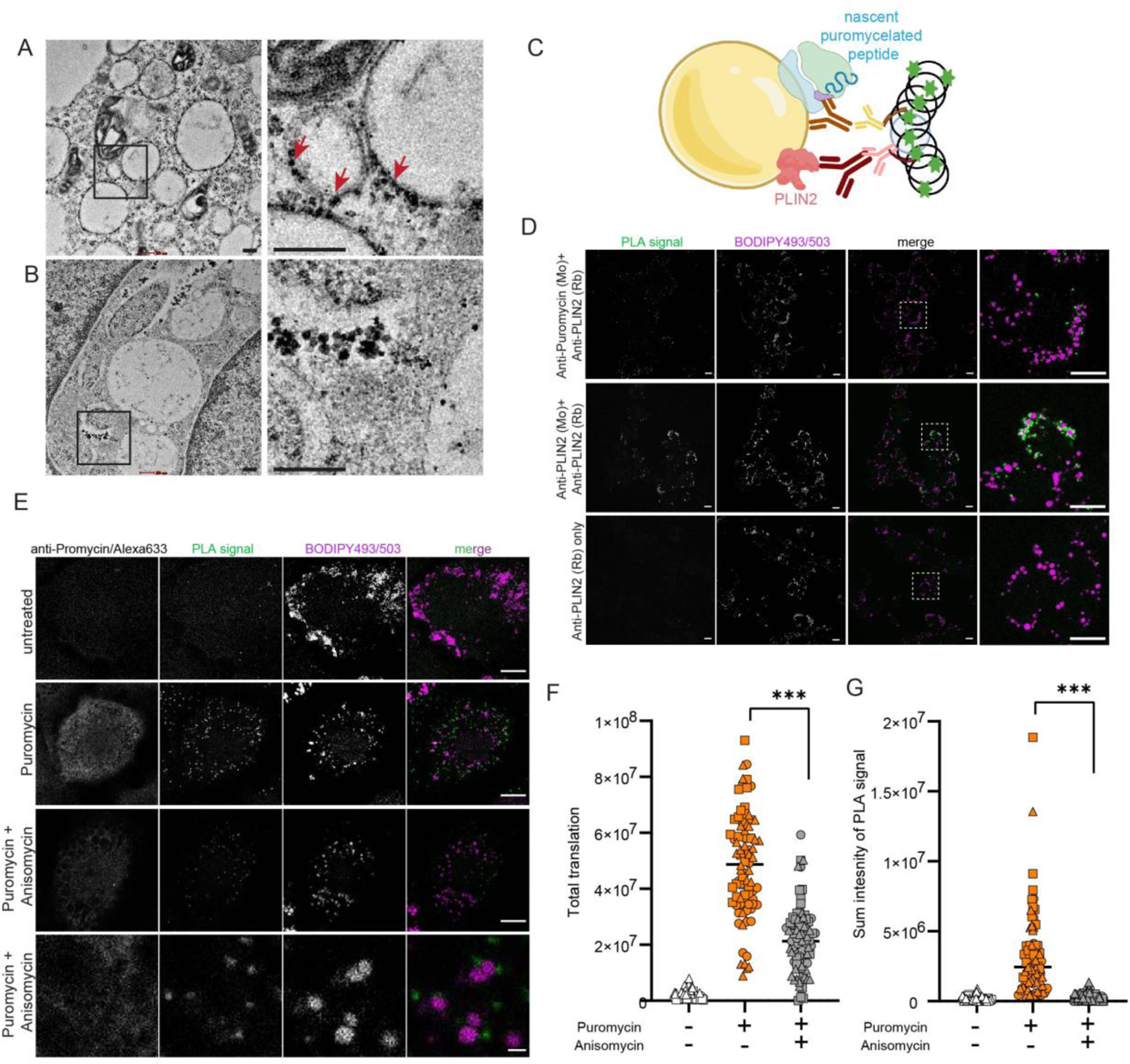
Functional ribosomes on lipid droplets: (A, B) Transmission electron microscopy (TEM) images of THP-1 cells showing lipid droplets (LDs) with and without ribosome association. Arrows indicate ribosomes attached to lipid droplets. Scale bar: 200 nm. (C) Schematic representation of the RPM-PLA assay derived LD proximal signal of functional ribosomes. (D) Proximity ligation assay (PLA) between PLIN2 and puromycylated peptides trapped in ribosomes was performed using antibodies against PLIN2 and puromycin. The PLA signal is displayed in monochrome and as a merged image in green, alongside the BODIPY 493/503 signal. Scale bar: 10 µm. (E) PLA between puromycylated peptides and PLIN2 following anisomycin treatment. The PLA signal is shown in monochrome and as a merged image in green with the BODIPY 493/503 signal. The anti-puromycin signal is also presented as a monochrome image. Scale bar: 10 µm in all except the lower-most panel where it is 1 µm. (F, G) Quantification of total translation was performed by measuring the total puromycin signal, while translational events on lipid droplets (LDs) were specifically assessed by quantifying the summed signal of PLA puncta. Data are presented on a per-cell basis, pooled from three independent experiments, with a minimum of 25 cells imaged per experiment. Symbols representing data from the same experiment are indicated with identical shapes. Statistical significance was determined using a two-tailed Mann-Whitney test (***p < 0.001).

Recently, stalled ribosomes have been described in neuronal cell lines. These retain puromycelated peptides even in the absence of emetine (Anadolu *et al*, 2024). However, these can be distinguished from actively translating ribosomes by the ability of anisomycin to compete with puromycin only in actively translating ribosomes. While puromycylation was significantly reduced upon Anisomycin treatment, it was not completely abrogated, suggesting that stalled ribosomes may indeed be present in the THP1 macrophages also (Figure 3E and F). To confirm whether the ribosomes detected on LDs are stalled, we performed RPM-PLA after treating cells with anisomycin. RPM-PLA signals dropped to background levels upon Anisomycin treatment, suggesting that ribosomes in proximity to LDs are not stalled (Figure 3E and G). Thus, these results collectively indicate that ribosomes are present near LDs and they are translationally active.

Due to conflicting reports claiming the limitations of emetine in retaining puromycylated peptides in active ribosomes (Hobson *et al*, 2020; Enam *et al*, 2020), we undertook a parallel approach to investigate LD-localized translation. We isolated LDs after puromycin treatment of oleic acid-loaded COS-7 cells, and immunostained them with anti-Puromycin antibody (Figure 4A). A ring of puromycin reactivity was present around lipid droplets further indicating translational activity in proximity of lipid droplets (Figure 4B). There was no signal of puromycelated peptides on LDs isolated from untreated cells indicating the specificity of the antibody (Figure 4C). This ring-like signal was not found when immunofluorescence was performed on LDs from untreated cells that were incubated with puromycin, confirming that puromycin itself was not accumulating on LDs (Figure 4C). To test the possibility of a cytosolic puromycelated peptide synthesized elsewhere but binding to lipid droplets during isolation, LDs were incubated with the post-nuclear supernatant of puromycin treated cells. The absence on anti-puromycin reactivity in this case confirmed that puromycylation of LDs was due to localized protein synthesis (Figure 4E).

**Figure 4:**
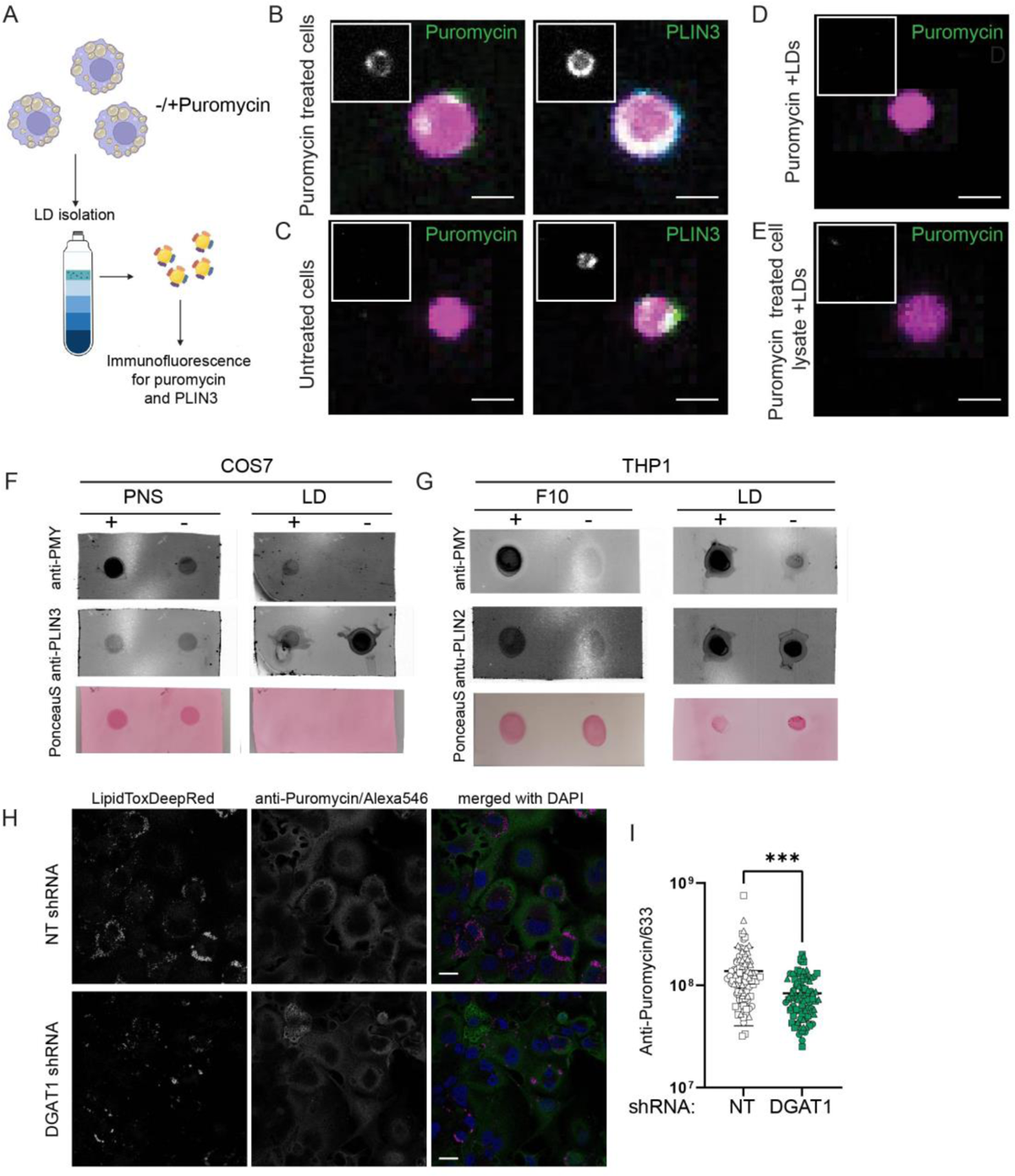
LDs contribute to localized and total translation (A) Schematic illustrating LD isolation and immunostaining using anti-puromycin and anti-PLIN3 antibodies following puromycin treatment. (B, C) Confocal images of lipid droplets from puromycin-treated and untreated cells, labelled with BODIPY 493/503 and immunostained for PLIN3 and anti-puromycin. The inset highlights the puromycin and PLIN3 signals surrounding the lipid droplets. Scale bar=1 µm. (D) Lipid droplets labelled with BODIPY 493/503 and immunostained with anti-puromycin antibody after *in vitro* incubation with puromycin. The inset shows the anti-puromycin signal surrounding the lipid droplets. (E) Lipid droplets labelled with BODIPY 493/503 and immunostained with anti-puromycin antibody following *in vitro* incubation with puromycylated peptides, with the inset highlighting the anti-puromycin signal around the lipid droplets. (F, G) Immunoblot against anti-Puromycin and anti-PLIN2/PLIN3 in both post-nuclear supernatant/F10 after ultracentrifugation and LD fraction from COS7 and THP1 cells. (H) Confocal images and signal quantification for anti-Puromycin signal of LipidTox DeepRed-stained THP1 macrophages stably expressing non-targeting (NT) or DGAT1-targeting shRNA. Data are on a per-cell basis, pooled from 3 independent experiments where a minimum of 25 cells were imaged per experiment. Symbols from the same experiment are represented by the same shapes. (****p < 0.0001, from a two-tailed Mann-Whitney test). Scale bar: 20 µm.

Puromycin is a global translational inhibitor (Aviner, 2020) and it could change the type of proteins or decrease the density of proteins present on lipid droplets. We argued that this could lead to random association of anti-PMY (puromycin) antibody at such “exposed” sites. To test this, the presence of puromycelated peptides in the LD fraction was verified by dot blot analysis (Figure 4F). Interestingly, puromycin treatment led to a decrease in PLIN3 levels following puromycin treatment (Figure 4F). LDs from THP1 cells were also stained with an anti-PMY antibody. However, for unknown reasons, the antibody exhibited non-specific binding to the isolated LDs despite specific binding in cells (Supplementary Figure 2, 3A and Figure 1E). This was also found to be the case in HEK293T cells (Supplementary Figure 3B). Despite this, a dot blot of proteins from the LD fractions indicated the presence of puromycelated peptides even in case of LDs isolated from THP1 macrophages. The levels of PLIN2 remained almost unchanged while total proteins in the LD fraction reduced with puromycin treatment, probably reflecting differential susceptibility of LD-localized proteins to translation inhibition (Figure 4G). Together, these data reveal that localized translation takes place at the LD and translation inhibition impacts differential recruitment of LD coat proteins. The latter observation motivated us to test if the presence of lipid droplets impacts global translation. Global puromycylation was used to assess translational activity. DGAT1 silenced THP1 macrophages exhibited a statistically significant decline in the total translation activity of the cell (Figure 4H and I), indicating a significant contribution made by the presence of LDs towards total cellular translation.

### Ribosome association with LDs is dynamically altered during infection with *M. tuberculosis*

After confirming the presence of translational activity on LDs we were interested to test if there would be some alteration upon infection, as a previous study has shown that several ribosomal proteins are increased in abundance in the LD fraction upon infection with *Mycobacterium tuberculosis* (Mtb) (Menon *et al*, 2019). We infected macrophages in the same manner as reported in this earlier study, with the virulent, H37Rv strain of Mtb. Transmission electron microscopy (EM) was used to test if the association of ribosome-like particles with LDs changed in response to infection. We classified this association into four categories. Class I associations were typified by LDs uniformly coated with ribosomes without a detectable ER membrane associated with it (Figure 5A). Class II associations were identified as ribosomes clustered or aggregated at some regions on the surface of the LD (Figure 5B). As LDs bud from the ER and can maintain a constant connection with the ER, we found some LDs in close proximity of ER and ribosomes were present at their contact sites. These were termed Class III associations (Figure 5C). However, the majority of LDs were found to be devoid of ribosomes (Figure 5D). These were classified as Class IV LDs. 18-25 fields with 285-585 LDs from a single experiment were analyzed and data from two independent experiments were analyzed. Across both experiments, we found an increase in Class II associations (clustered ribosomes) and a decrease in ribosome-free LDs in the case of infection (Figure 5E). This could explain earlier findings of higher ribosomal protein content in the LD fraction of Mtb infected macrophages compared to uninfected macrophages (Menon *et al*, 2019).

**Figure 5:**
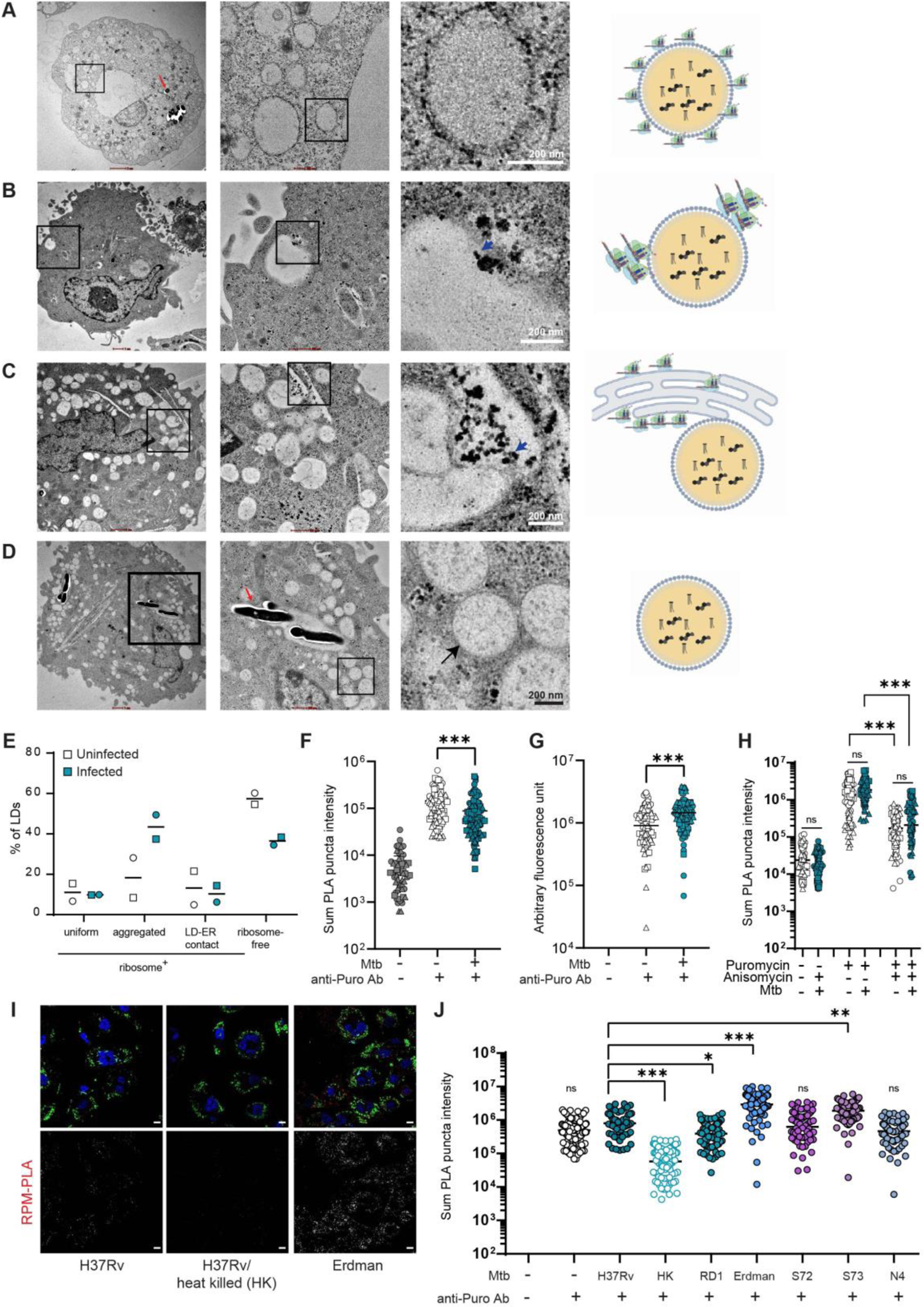
Alteration in translation activity on lipid droplets following infection (A-D) TEM images depicting various types of associations between lipid droplets and ribosomes in both infected and uninfected macrophages. From left to right: individual cells, magnified regions, and digitally magnified individual LDs. The red arrow points to Mtb within the cells, while the blue arrow marks ribosomes surrounding the lipid droplets, and the black arrow indicates a ribosome free-LD. (A) Uniformly coated with ribosomes (B) Ribosomes aggregated on LDs (C) ribosomes are present because of proximity of ER membrane and (D) ribosome free LDs. (E) Quantification of the different types of associations between ribosomes and lipid droplets in infected and uninfected macrophages. Data represent the percentage of these interactions from 15-20 cells per experiment (N=2). (F, G) Quantification of the summed signal from PLA puncta and anti-puromycin signal in infected and uninfected cells. Anti-Puromycin-Alexa647 was used here. Data are presented on a per-cell basis, pooled from three independent experiments, with a minimum of 25 cells imaged per experiment. Symbols representing data from the same experiment are indicated by identical shapes. Statistical significance was determined using a two-tailed Mann-Whitney test (***p < 0.001). (H) Sum RPM-PLA signal where cells were treatment with Anisomycin or untreated control prior to Puromycin treatment. Data are presented on a per-cell basis, pooled from three independent experiments, with a minimum of 25 cells imaged per experiment. Symbols representing data from the same experiment are indicated by identical shapes. Statistical significance was determined using a two-tailed Mann-Whitney test (***p < 0.001). (I) Representative confocal images from infected macrophages (Mtb:cyan; LD: green, PLA:red, DAPI:blue), Scale bar=10 µm. (J) Sum RPM-PLA intensity in THP1 macrophages infected with indicated strains of *M. tuberculosis*. Data are from a single experiment. Statistical significance was determined using Kruskal Wallis test (***p < 0.001, **p<0.01, *p<0.05, ns=not significant).

We performed RPM-PLA and found a decrease in the total signal of PLA puncta upon infection (Figure 5F), despite higher levels of total puromycylation (Figure 5G). RPM-PLA works if there is a distance of 40 nm between two antigens. In a cluster of ribosomes as seen in TEM, we suspect that there would be an increased distance between peripheral ribosomes and lipid droplets and hence there would be less likelihood of PLA to happen, potentially decreasing the signal. This made us question whether an increase in clustering of ribosomes was due to stalled or active ribosomes. To test this, we again did PLA after treating cells with anisomycin followed by puromycin. From these results, it could be seen that the number of PLA puncta decreased to similar extent in both infected and uninfected conditions which further indicates that ribosomes clustered in case of Mtb infection are not stalled (Supplementary Figure 5). We noticed that the comparison of uninfected and infected condition was sensitive to the type of anti-puromycin antibody we used in the RPM-PLA assay; while the usage of the Alexa fluor tagged antibody revealed a small but statistically significant difference between uninfected and infected cells (Figure 5F), the usage of an untagged antibody revealed no difference between the two conditions (Figure 5H). This small difference is also likely due to sensitivity of the assay to placement of the ribosomes relative to the surface of the LDs.

Previously, Mtb induced changes in the ribosomal protein content of LDs were observed in comparison to uninfected macrophages and macrophages infected with heat killed Mtb. We sought to investigate this further with the RPM-PLA assay. While the RPM-PLA reactivity of uninfected and live Mtb infected macrophages was comparable, we found a larger decrease in the RPM-PLA reactivity of macrophages infected with heat killed Mtb. To further understand if the LD-associated ribosome activity is virulence related, we compared the RPM-PLA reactivity across various strains of Mtb. Mtb actively alters the endocytic machinery and the LD proteome. It’s ability to exit the phagolysosome and escape into the cytosol is a major virulence feature attributed to a genomic locus known as *region of difference 1 (RD1)* (Lewis *et al*, 2003; Simeone *et al*, 2015). The ΔRD1 strain exhibited a small but statistically significant reduction in the RPM-PLA signal. This suggested that the ability of Mtb to activity manipulate the host cell plays a role in LD-associated ribosomal activity. Another virulent strain of Mtb, the Erdman strain, led to a significantly higher LD-associated ribosomal activity. While both H37Rv and Erdman belong to the same lineage (Lineage 4), we sought to compare the RPM-PLA signal induced by other pathogenic clinical strains. The ancient lineage (Lineage 1) strains of Mtb have previously been shown to be poor inducers of the Type I IFN response and shown to reside poorly in the lysosomal compartment while the modern lineage prevalent in India belongs to Lineage 3 and is more pathogenic (Shankaran *et al*, 2022). We compared the LD-associated ribosomal activity across all these strains and found that one of the lineage 1 strains, S73 exhibited higher activity in comparison with H37Rv while the other strain of the same lineage did not. In addition, the strain N4 of Lineage 3 did not exhibit differences in LD-associated ribosome activity. Together, these results revealed that LD-associated ribosomal activity is regulated in a stimulus dependent manner in macrophages, and likely offers a nuanced means of impacting host responses to infection.

## Discussion

Till date it was known that LD proteins are either recruited from the cytosol or imported from ER (Olarte *et al*, 2022). In this study we provide evidence that LDs in mammalian cells harbor intact and functional ribosomes on their surface. This opens the possibility of a new mechanism of targeting newly synthesized proteins to the LD. Our attempts to identify transcripts of PLIN2, the major LD coat protein in THP1 macrophages was unsuccessful (data not shown), likely because proteins such as PLIN2 and PLIN3 are indeed exchangeable or CYTOLD proteins, relying on cytosolic ribosomes for synthesis (Bickel *et al*, 2009; Bulankina *et al*, 2009; Kimmel & Sztalryd, 2016). PLIN1, another LD coat protein which has recently been described to be an ERTOLD protein, is not expressed in THP1 macrophages (Majchrzak *et al*, 2024; Menon *et al*, 2019; Mejhert *et al*, 2020). Several canonical ERTOLD proteins rely on either the Signal recognition protein (SRP) and its receptors (SR), the Sec61-dependent or the ER-membrane complex dependent co-translation pathways for ER localization (Leznicki *et al*, 2022; Abell *et al*, 2002; Monier *et al*, 1995). In addition some LD-localized proteins such as UBXD8 are delivered to the ER in a post-translational manner, in a Pex19/Pex3 dependent manner, wherein Pex3 serves as an ER-membrane localized receptor for UBXD8 (Yamamoto & Sakisaka, 2018; Schrul & Kopito, 2016). We expect that for high abundance proteins to be synthesized at the LD, we would expect to find a much higher density of ribosomes. Our TEM data suggested that ribosomes are less likely to decorate the entire surface of LDs. Immunofluorescence data revealed different densities and staining patterns of the individual ribosomal protein RPS9 and RPL4 on isolated LDs. This suggests that LDs are heterogenous in their protein synthesis capacity.

How could ribosomes be recruited to LD? It is possible that the recruitment of functional ribosomes requires a receptor protein on LDs. Alternatively, ribosomes could be recruited in a transcript or product protein dependent manner. While the former would require the presence of RNA-binding proteins on the LD, the latter would require translation of proteins that have an inherent capacity to bind to LDs. We and others have reported several RNA binding proteins in LD proteomes; it is possible that some of these are involved in recruiting RNA for translation to the LD (Menon *et al*, 2019; Cho *et al*, 2006; Mejhert *et al*, 2020; Rösch *et al*, 2016). Our data also supports the latter possibility as we found puromycylated proteins localized to LD only when the latter were isolated from puromycin treated cells but not lysates from puromycin treated cells were incubated with them. This confirmed that what our assays captured were newly synthesized proteins being made on LDs within cells. Identification of RNA localized to LDs in the future will help in understanding if one or more of these models are likely to be valid.

Although there have been reports suggesting the presence of RNA within the lipid droplet core, the hydrophobic nature of the core makes it unlikely for RNA to be present inside it (Dvorak *et al*, 2003). Recently, a human long non-coding RNA, Lipid-Droplet Transporter (*LIPTER*), was shown to be associated with lipid droplets (Han *et al*, 2023). While RNA-FISH have demonstrated colocalization of *LIPTER* and LDs, biochemical evidence confirming the presence of RNA on lipid droplets falls short. While our study does not provide sufficient evidence to assert that rRNAs are specifically bound to LDs, it is the first to demonstrate the presence of RNA within the LD fraction. Our data does however support the likelihood of rRNA being present in the LD fraction in the context of a complete ribosome. Based on our findings, we also caution that simple flotation-based isolation of LDs should not be used to identify LD-associated transcripts.

Many early ultrastructure-based studies on lipid droplets were conducted on cells of the innate immune system during infection conditions (D’Avila *et al*, 2006; Melo *et al*, 2003; Pacheco *et al*, 2002). A greater molecular understanding of the nature of lipid droplets re-ignited an interest in how pathogens manipulate these organelles. In case of tuberculosis, the oldest documented infectious disease that is still the number one reason for death due to a single pathogenic organism [Global Tuberculosis Report 2024], the presence of lipid droplet rich macrophages are a hallmark feature of the pathology (Jaisinghani *et al*, 2018; Dawa *et al*, 2021; Kim *et al*, 2010). Furthermore, the LD proteome of macrophages following *Mtb* infection undergoes significant changes, with most notably an increase in the abundance of several ribosomal proteins and other factors associated with protein synthesis (Menon *et al*, 2023). In our study, we used electron microscopy and RPM-PLA to show that infection with the virulent strain H37Rv significantly alters LD associated translation. We found increased clustering of ribosomes on lipid droplets compared to uninfected cells, suggesting an increased density of ribosomes on mRNAs. Importantly, the frequency of such associations increases while the frequency of ribosome-free LDs decreases with infection, pointing towards an increased LD-associated translation upon infection. It is known that ribosomal density on mRNA regulates translation; insufficient ribosomes result in low translation rates, while excess ribosomes can cause traffic jams on mRNAs reducing translation efficiency. Studies have shown that an optimal ribosome density is necessary for maximum translation efficiency, with mathematical modeling suggesting optimal density is about half of the maximum possible (Zarai *et al*, 2016). Moreover, ribosome density has also been correlated with length of the ORF (Arava *et al*, 2003). Therefore, translation control could be one of the reasons for the altered LD associated translation after Mtb infection and the clustering of ribosomes during Mtb infection could indicate the presence of different mRNAs pools near lipid droplets. Control of translation is a host response to many viral infections, induced as part of the Type-I IFN response. That this largely anti-viral response, is also induced upon bacterial infection, is now an established paradigm. The Type-I IFN response is a pathogenic immune response in case of tuberculosis, with high expression being correlated with active infection in humans and disease susceptibility in mice (Kotov *et al*, 2023; Berry *et al*, 2010). Inhibition of the response of macrophages to Type I IFN signaling has emerged as a means for targeting bacterial growth and disease susceptibility (Shankaran *et al*, 2023; Dorhoi *et al*, 2014). Protein Kinase R (PKR) is activated during this signaling (Smyth *et al*, 2020). PKR is a double stranded RNA-activated, serine/threonine kinase (Zhang *et al*, 2001), which phosphorylates the translation initiation factor EIF2α thereby inhibiting translation. While human macrophages lacking PKR exhibit lack of bacterial control, the effect of PKR inhibition in mouse macrophages and mouse models of tuberculosis is debated (Wu *et al*, 2012; Mundhra *et al*, 2018; Smyth *et al*, 2020). PKR activation to limit viral translation while still enabling key host defense proteins to be translated seems like an evolutionary advantage to limit viral growth, the regulation of host translation in case of bacterial infections is unexplored. The observation that dead bacilli suppressed LD-associated ribosomal activity indicates that the ability of LDs to partake a role in cellular translation is very much dependent on its ability to sense a foreign agent. Indeed, the inability to make LDs by silencing DGAT1 leads to an attenuated pro-inflammatory response by macrophages to the bacterial endotoxin lipopolysaccharide and live Mtb (Jaisinghani *et al*, 2018; Castoldi *et al*, 2020). Future studies in the area of LD-localized translation will help understand how LDs play such a central role in host defense.

Our study provides evidence that lipid droplet-associated ribosomes are active and synthesizing proteins at the lipid droplet. However, one limitation is that we do not yet have a comprehensive list of transcripts translated by these ribosomes. The relatively low amounts of RNA isolated from gradient purified lipid droplets, possible heterogeneity in RNA present on individual lipid droplets, and the dynamic nature of the translation complex has precluded our ability to assess this at this point. However, this study paves the way to understand a potentially pivotal role played by localized translation on LDs, particularly in host defense.

## Materials and Methods

### Key reagents

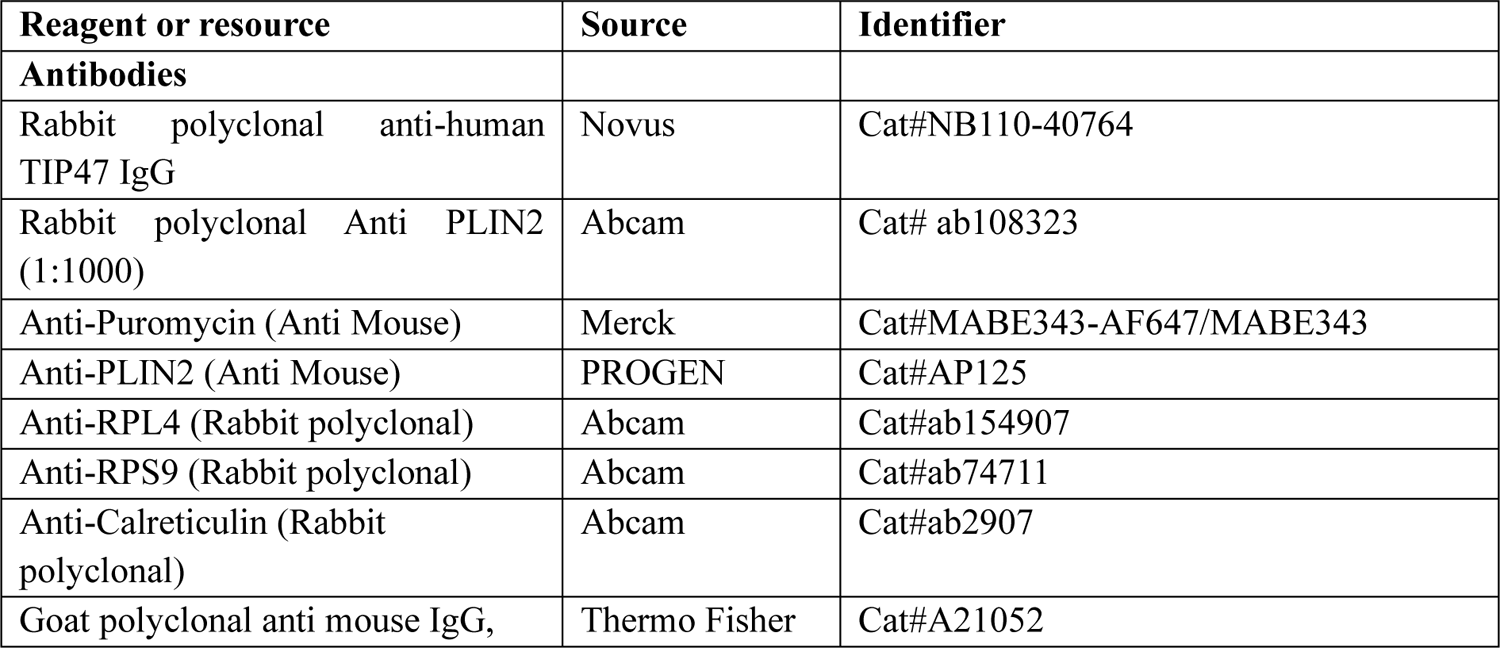
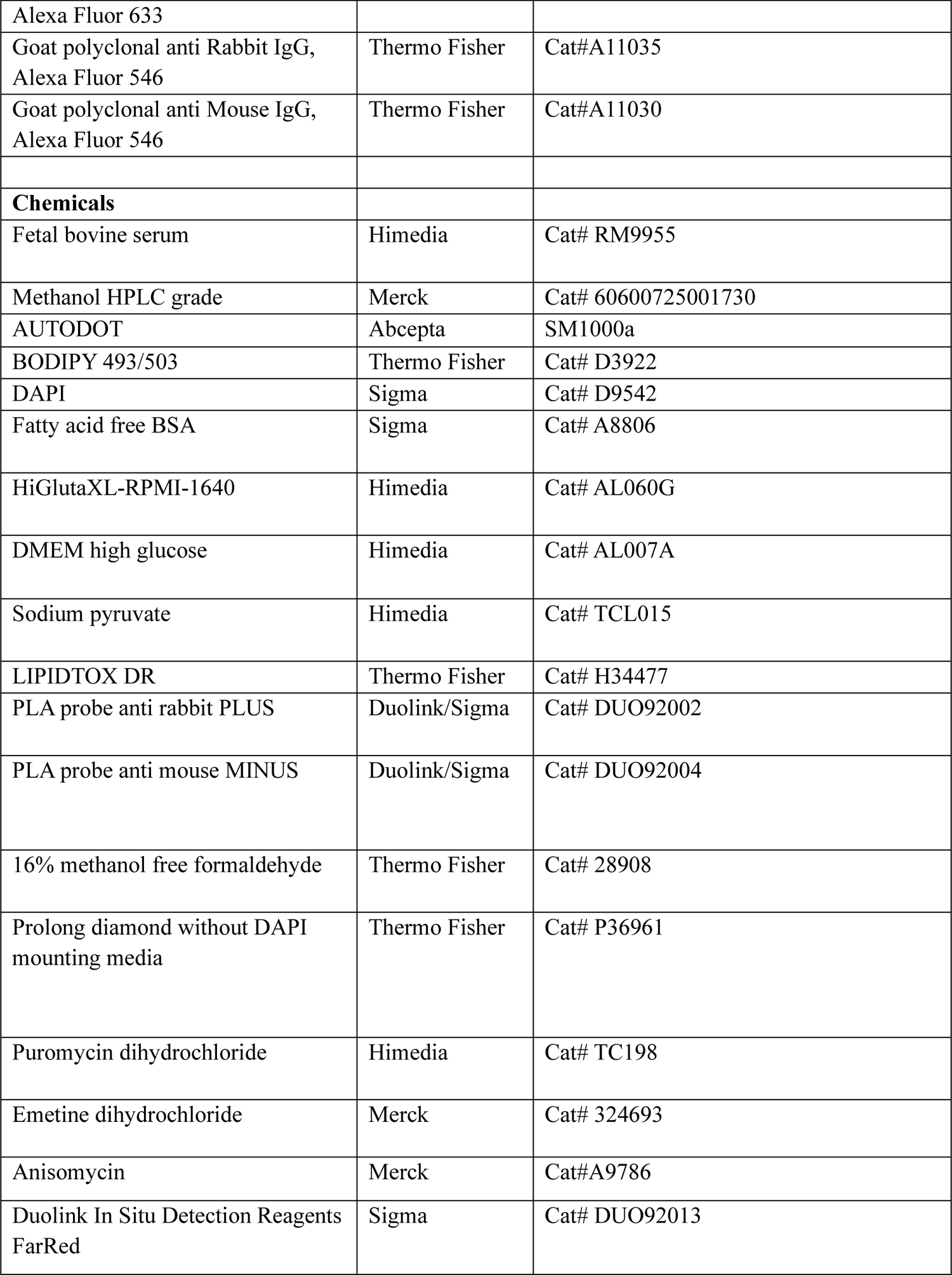
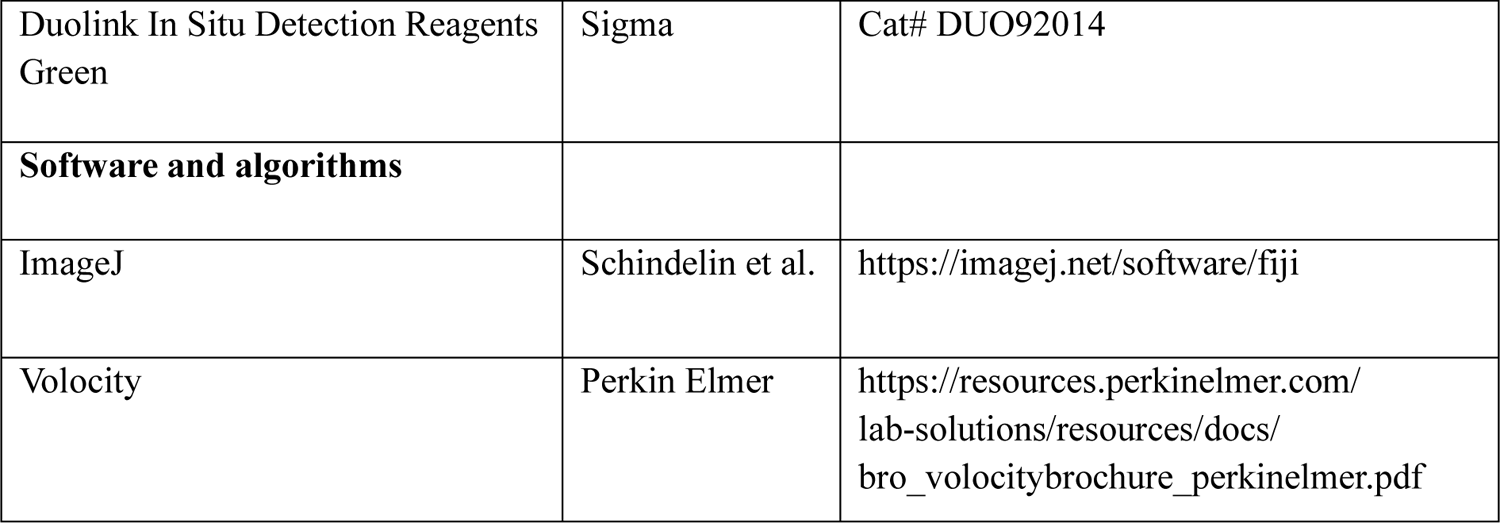

### Cell culture

THP-1 monocytes, purchased from ECACC, were used in this study. These cells were cultured in RPMI 1640 medium supplemented with 10% FBS and 1 mM sodium pyruvate. THP-1 monocytes were differentiated by treatment with 100 nM Phorbol-13-myristate 13-acetate (PMA) for 24 hours, followed by a media change. Experiments were conducted 2 days after this media change unless otherwise specified. COS-7 cells, used for LD isolation, were cultured in DMEM high-glucose medium. For LD isolation from COS7 cells, media was changed and fresh media containing 200 mM of oleic acid was added for 48h.

### Infection

*Mycobacterium tuberculosis* (Mtb) strains expressing mCherry were cultured to an OD600 of 0.4–0.6 in 7H9 media supplemented with Middlebrook ADS, supplemented with 0.5% glycerol and 0.05% TWEEN 80, and hygromycin at a concentration of 50 μg/ml. The cultures were harvested by centrifugation at 3000 rpm for 5 minutes using a swing-out bucket rotor, washed twice with phosphate-buffered saline (PBS) containing 0.05% Tween 80, and resuspended in PBS. To prepare a single-cell suspension (SCS), the bacterial suspension was centrifuged at 800 rpm for 12 minutes in a swing-out bucket rotor. The supernatant was carefully transferred to a new tube using a pipette and treated as the SCS. The optical density of the SCS was measured at 600 nm, and bacterial density was calculated using the conversion factor of 1 OD = 5 × 10⁸ bacilli/mL. Macrophages were infected with Mtb at a multiplicity of infection (MOI) of 3 for 16 hours, using half the volume of media typically used for macrophage culture.

### LD isolation for EDTA and puromycin experiments

Cells were gently scraped in media and then centrifuged in a fixed angle rotor for 10 min at 500 g for 15 minutes at 4^0^C. The cell pellet was washed twice with chilled PBS and then resuspended in 2 mL lysis buffer A (10 mM Tris, 1 mM EDTA, 1 mM EGTA, 0.25 M Sucrose and 100 mM KCl). The lysis of cells was completed using a 3 mL Dounce homogenizer (40 strokes/sample). The total cell lysate was centrifuged at 900 g for 10 min at 4^0^C. The supernatant so obtained was treated as the post-nuclear supernatant (PNS). An aliquot of PNS was kept for RNA isolation.

Gradients were prepared and resolved in a BECKMAN Ultra-Clear tube (Cat #344061). The PNS was layered between a sucrose cushion (1 mL of buffer B (20 mM Tris, 1 mM EDTA, 1 mM EGTA, 1.08 M Sucrose and 100 mM KCl, pH7.4)) and sucrose gradient: 2 mL of buffer C (20 mM Tris, 1 mM EDTA, 1 mM EGTA, 0.20 M Sucrose and 100 mM KCl), 2 mL of buffer D-20 mM Tris, 1 mM EDTA, 1 mM EGTA, 0.13 M Sucrose and 100 mM KCl, followed by 4 mL of Buffer E-25 mM Tris, 1 mM EDTA, 1 mM EGTA on the top. All the buffers used contained protease inhibitor cocktail (Roche). The tubes were centrifuged at 1,86,700 g in an SW32.1 Ti rotor in a Beckman Coulter Optima XPN-100 ultracentrifuge. One millilitre fraction from the top were collected.

### LD isolation from 15 flasks and DGAT1 KD experiments

Cells were gently scraped in media and then centrifuged in a swing out bucket rotor at 500 g for 15 min. The cell pellet was washed twice in PBS and then resuspended in 2.5 mL of lysis buffer A’ (20 mM Tris, 1 mM EDTA, 1 mM EGTA, 100 mM KCl buffer (pH 7.4)), followed by incubation on ice for 5 min. The lysis of cells was completed using a Dounce homogenizer (40 strokes/samples). The cell lysate was collected in 15 mL conical tubes to which 2.5 mL of Buffer B’ (20 mM Tris, 1 mM EDTA, 1 mM EGTA, 100 mM KCl buffer, 1.08 M sucrose (pH 7.4)) was added and mixed. This was treated as the total cell lysate. The total cell lysate was centrifuged at 900 g for 10 min. The supernatant so obtained was treated as the post-nuclear supernatant (PNS). An aliquot of PNS was kept for RNA extraction.

Gradients were prepared and resolved in a BECKMAN Ultra-Clear tube (Cat #344061)/ Thinwall polypropylene tube (Cat #331372). The PNS was layered between a sucrose cushion (1 mL of buffer B (20 mM Tris, 1 mM EDTA, 1 mM EGTA, 1.08 M Sucrose and 100 mM KCl, pH7.4) and sucrose gradient (2 mL of buffer C (20 mM Tris,1 mM EDTA, 1 mM EGTA, 100 mM KCl buffer, 0.27 M sucrose (pH 7.4), 2 mL of buffer D-20 mM Tris, 1 mM EDTA, 1 mM EGTA, 0.13 M Sucrose and 100 mM KCl, followed by 1 mL of Buffer E-25 mM Tris, 1 mM EDTA, 1 mM EGTA on the top). All the buffers used contained protease inhibitor cocktail (Roche). The tubes were centrifuged at 1,86,700 g in an SW32.1 Ti/SW 41Ti rotor in a Beckman Coulter Optima XPN-100 ultracentrifuge. One millilitre fraction from the top were collected.

### RNA isolation and qRT-PCR analysis

Total RNA extraction from the PNS and LD fractions was performed using RNAzol RT reagent (Sigma-Aldrich, R4533) following the manufacturer’s protocol for liquid samples. The RNA was eluted in 50 µl of RNase-free water. cDNA was synthesized from 12 µl of RNA from the LD fraction and 5.5 µl of RNA from the PNS fraction using random hexamers. (For reference, 5 ml of PNS was loaded onto the gradient for LD isolation, with a total gradient volume of 11 ml. The ratio between the total gradient and PNS is 11/5 = 2.2. Therefore, if 12 µl of RNA from the LD fraction was used, the corresponding RNA volume from the PNS fraction would be 11/2.2 = 5.45 µl).

cDNA synthesis was carried out using the Verso cDNA synthesis kit (Thermo Fisher Scientific, Cat. No AB1453B) with random hexamers. The resulting cDNA was diluted 5-fold, and 2 µl of the diluted cDNA was used for gene expression analysis. qRT-PCR (Quantitative real time PCR was performed on Light cycler 480 (Roche) instrument with KAPA SYBR Fast Universal qPCR master mix (Cat. No SFLCKB, Roche). Transcripts levels were normalized to dilution factors for quantitation in PNS and LD fraction. The following primers were used in the study:

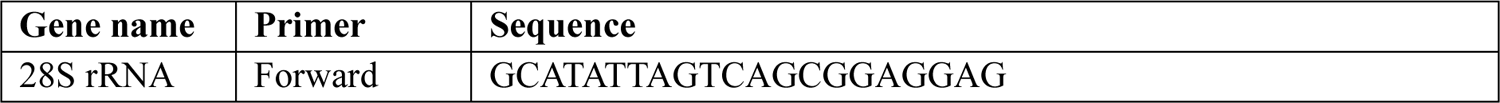
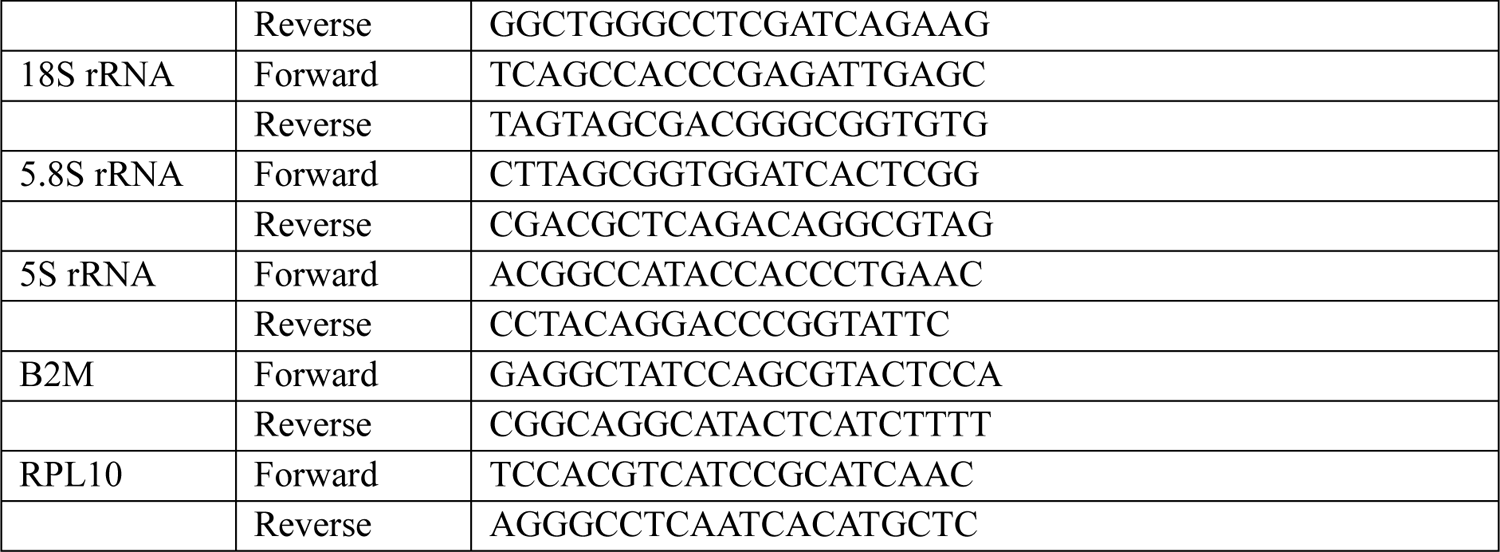

### Immunostaining

THP1 macrophages were given three quick washes with PBS before fixation with 4% methanol-free formaldehyde in 1X PHEM buffer (PIPES: 60 mM, HEPES: 27.28 mM, EGTA: 10 mM, MgSO₄: 4.02 mM, pH = 7). Permeabilization was done using 0.5% saponin in 1X PHEM buffer for 10 minutes at room temperature (∼25°C) on a shaker. Following permeabilization, three quick washes with 1X PBS were given and blocking was done using 3% BSA in 1X PHEM buffer + 0.05% Tween 20 for 1h at RT. Primary antibody incubation was performed overnight in blocking buffer at 4ᣞC. Subsequently, coverslips were washed thrice for 5 min each with washing buffer (0.1% BSA in 1X PBS + 0.05% Tween 20), followed by incubation with secondary antibody solution (1% BSA in 1XPHEM with 0.05% Tween 20) for an hour on a shaker at room temperature. Samples were further washed thrice, 5 min each in washing buffer and subsequently stained with BODIPY 493/503 and DAPI for 30 min. Coverslips were further mounted on glass slides before visualizing using the Leica SP8 laser scanning confocal microscope.

### RiboPuromycylation Method

THP1 cells were seeded on glass coverslips in a 24 well plate. RPM was performed as described by David et al., 2012. Depending on the experiments, cells were pretreated with different inhibitors. Cells were pretreated with 115.14 µg/mL emetine for 15 minutes in media before addition of 49.54 µg/mL puromycin for further 15 minutes. For stalled ribosomes experiments, cells were pretreated with 2.49 µg/mL of Anisomycin for 15 minutes before addition of 49.54 µg/mL of puromycin for further 15 minutes. Immunostaining of cells were performed after permeabilization and fixation as described in the previous section using anti-puromycin antibody (1:250 dilution). Either anti-Puromycin-Alex647 or anti-Puromycin (untagged) antibody was used for the ribopuromycylation.

### Proximity ligation assay

1,50,000 THP-1 cells were seeded in each well of a 24-well plate. After treatment with translation inhibitors (as described in the RPM method section), the cells were washed three times with 1X PBS and fixed using 4% methanol-free formaldehyde in PHEM buffer for 30 minutes. Permeabilization was performed with 0.5% saponin in PHEM buffer for 10 minutes at room temperature on a shaker. The cells were then washed three times with 1X PBS and blocked using the blocking solution provided in the DUOLINK PLA kit on parafilm in a humid chamber for 1 hour at 37°C. Primary antibody incubation was carried out overnight at 4°C in 3% BSA diluted in PHEM buffer with 0.05% Tween 20. The next day, the cells were washed twice with buffer-A (provided in the kit) for 5 minutes each and incubated with the plus and minus PLA probes for 1 hour at 37°C. After washing twice with buffer-A, the probes were ligated using the ligase provided in the kit at 37°C for 30 minutes. The PLA signal was then amplified with the amplification reagents from the kit for 100 minutes at 37°C. The cells were washed twice with buffer-B for 10 minutes each, stained with BODIPY 493/503, and mounted on glass slides for observation under a laser scanning confocal microscope (LSCM).

### Immunostaining of purified lipid droplets

Lipid droplets were isolated from COS-7 or THP-1 cells using ultracentrifugation, with or without treatment with 49.54 µg/mL of puromycin for 30 minutes. After centrifugation, the top 1 ml of the density gradient fraction was collected. For staining, 50 µl of this fraction was mixed with 50 µl of a primary antibody solution (1:100 dilution) prepared in 6% BSA in 1X PBS and incubated for 1 hour at room temperature (RT). Following the incubation, 100 µl of 1X PBS was added, and the samples were centrifuged at 15,000 g for 10 minutes. Then, 50 µl of the supernatant was collected and mixed with 50 µl of GAR-546 (1:300 dilution) in 6% BSA in 1X PBS. The samples were incubated again at RT for 1 hour, after which 100 µl of 1X PBS was added, and centrifugation at 15,000 g for 10 minutes was repeated. Finally, 50 µl of the supernatant was collected and stained with BODIPY 493/503 before imaging using confocal microscopy.

To assess puromycin accumulation in lipid droplets (LDs), LDs isolated from untreated cells were incubated with 49.54 µg/mL puromycin in a test tube at 37°C for 30 minutes. After the incubation, the LDs were stained and observed under a microscope.

In a separate experiment, puromycylated peptides were added to LDs isolated from untreated cells, and the mixture was incubated at 37°C for 30 minutes. Following the incubation, these LDs were stained and imaged using confocal microscopy. To obtain puromycylated peptides, cells were treated with puromycin as described above and processed for isolation of the PNS fraction. This was used as a source of puromycylated peptides. The ratio of LD fraction used relative to the total gradient is 50 µl/11,000 µl = 1/220. Thus, the corresponding volume of PNS (post-nuclear supernatant) from puromycin treated cells to be added to LD fraction was calculated as 2000 μl×1/220=9.09 μl. LDs with puromycylated peptides were incubated in a test tube at 37°C for 30 minutes. After the incubation, the LDs were stained and observed under a microscope.

### Protein dot blot

For delipidation of samples from PNS and LD fraction, samples were mixed with 4 volumes of ice-cold acetone and incubated overnight at −80^0^C, followed by centrifugation at 15,500 g for 15 min. The pellet so obtained was washed with acetone. The protein pellets were air-dried and then resuspended in 50 µl of 50 mM triethylammonium bicarbonate buffer (TEABC; pH 8.0) containing 0.1% SDS. Immunoblotting for puromycylated peptides and lipid droplets coat proteins was performed.

### Transmission electron microscopy

TEM was performed as previously described (Choudhary *et al*, 2015). THP1 cells were washed with 0.1 M phosphate buffer to get rid of cell culture media. Briefly, cells were fixed with primary fixative (2.5% glutaraldehyde, 1.25% paraformaldehyde, and 0.1 M phosphate buffer, pH 7.0) for 15 min at RT. Cells were further resuspended in 1 ml fresh fixative and incubated on ice for 1 hour. Cells were washed with 0.1 M phosphate buffer followed by secondary fixation with 1% osmium tetroxide (OsO4) in 0.1 M phosphate buffer for 1 hour at RT followed by washing with distilled water twice. Next, cells were incubated with 1% uranyl acetate for 1 hour at RT. Subsequently, samples were dehydrated by incubating in increasing concentrations of ethanol (30%, 50%, 70%, 80%, 90%, 95%, and 100%) for 10 minutes each, with two more incubations in 100% ethanol. The samples were subsequently embedded stepwise using Spurr’s low-viscosity resin. Samples were infiltrated for 2 h each with a 3:1, 1:1, 1:3 ethanol / embedding media mixture. Cells were incubated overnight with 100% fresh resin. The next day, cells were again resuspended in fresh 100% resin for 2–3 h, transferred into BEEM capsules (EMS), and cured at 70°C for 4 days. Semi and ultrathin sections were obtained with a Diamond knife (Diatome) on an ultramicrotome (Ultracut UCT; Leica Microsystems), collected on 200 mesh copper grids (EMS), poststained with uranyl acetate and lead citrate, and visualized with a Talos S200 transmission electron microscope (TEM; Thermo Fisher Scientific), operating at 200 kV. Pictures were recorded on a below mounted 4k × 4k BM-Ceta (CMOS) camera.

### Quantification and statistical analysis Image acquisition and analysis

All images of COS-7 cells, THP-1 cells, and purified LDs were captured using the Leica SP8 laser scanning confocal microscope, with a pixel size of 90 nm x 90 nm. Post-acquisition image analysis was performed using Volocity image analysis software.

### Quantification of sum PLA and total puromycylation reaction

Single confocal plane images of each sample were acquired. Post-acquisition, images were analysed using Volocity image analysis software. Object segmentation was conducted by setting an intensity bandwidth and defining minimum object size for each experiment. After this sum PLA puncta per cell was measured by drawing a ROI around each cell using bright-field images as a reference.

Total puromycylation signal was determined by measuring the integrated intensity of the anti-puromycin signal, with ROIs defined around cell perimeters using bright-field images as a reference.

## Statistics and analysis

Statistical analyses were conducted using GraphPad Prism. The normality of data was assessed using the Shapiro-Wilk normality test. For non-normal distributions, a non-parametric Kruskal Wallis or Mann-Whitney test was applied, while for normal distributions, a parametric Student’s t-test was used. All illustrative images were generated using BioRender.

## Acknowledgements

This work was funded by Swarnajayanti Fellowship Research Grant support from DST and SERB awarded to SG (DST/SJF/LSA-01/2018-19 and SB/SJF/2019-20/03). SG also acknowledges funding support from CSIR OLP2306 and STS0021 for facilities crucial to this work. Research in VC lab is supported by an Indo Swiss Joint Research Programme of Ministry of Science & Technology, Department of Biotechnology, Government of India (IC-12044(11)/6/2021-ICD-DBT, an Early Career Intramural Project of the All India Institute of Medical Sciences (AIIMS), New Delhi (A-1012/2023/RS), and DBT/Wellcome Trust India Alliance Fellowship (Grant IA/I/20/2/505191). RJ acknowledges Department of Biotechnology for junior and senior research fellowship awarded to him. AN acknowledges University Grants Commission for her fellowship. DM acknowledges CSIR for Senior Research Fellowship awarded to him. We acknowledge the central instrumentation facility University of Delhi South Campus for assistance with ultracentrifugation in some of the experiments.

## Author contributions

Conceptualization: SG; Data Analysis: RJ and SG; Funding acquisition: SG; Investigation: RJ, DM, AN, AK; Methodology: RJ, DM, VC, SG; Project Administration: SG; Resources: VC and SG; Supervision: SG; Visualization: RJ and SG; Writing: RJ and SG; Writing-Review & Editing: RJ, DM, AN, AK, and SG.

---

**Supplementary Figure 1.**
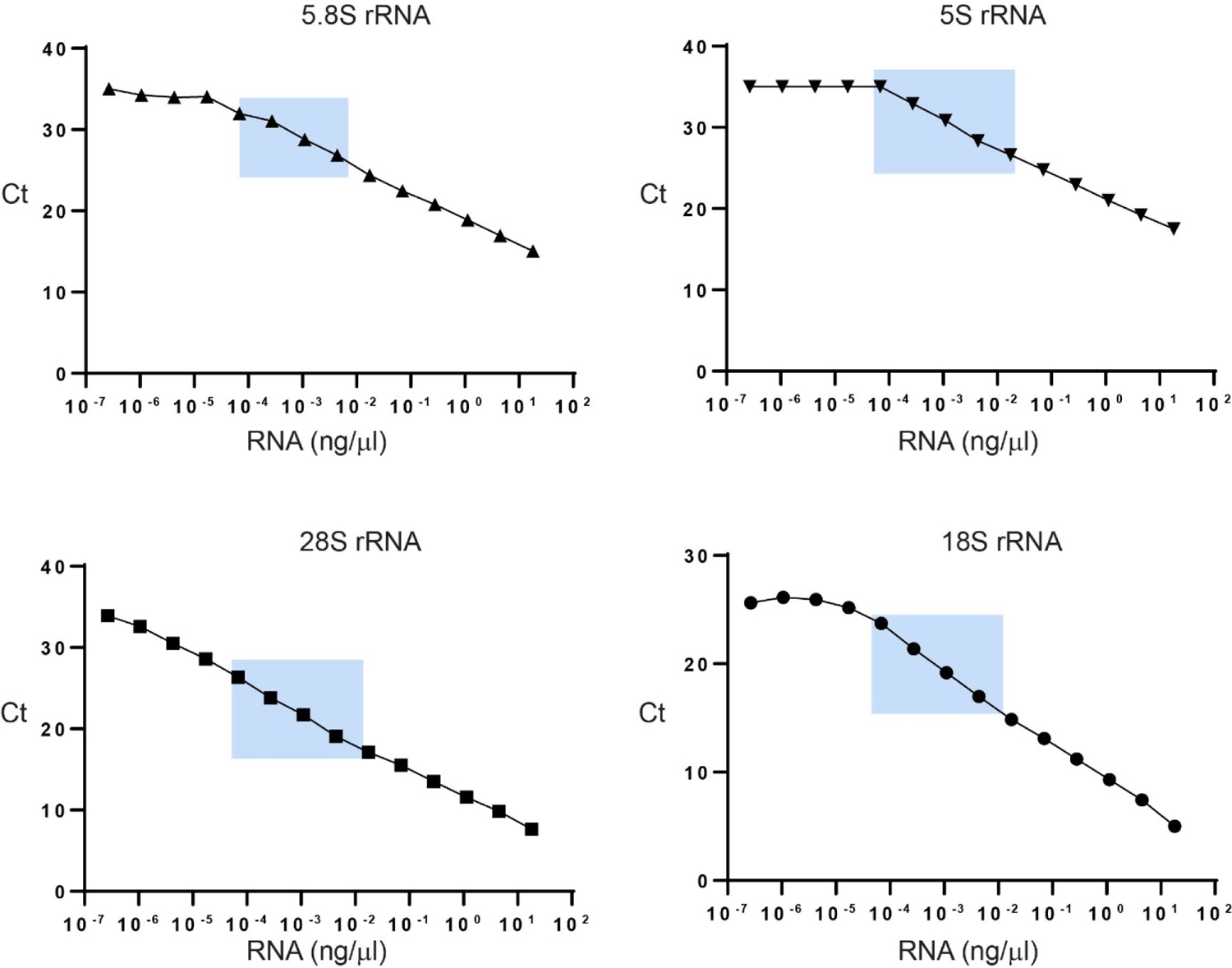
Ct values obtained for a dilution series of cDNA corresponding to RNA concentration. Boxed in blue color are the Ct values obtained for the LD fraction when using 1 T75 flask for LD isolation.

**Supplementary Figure 2.**
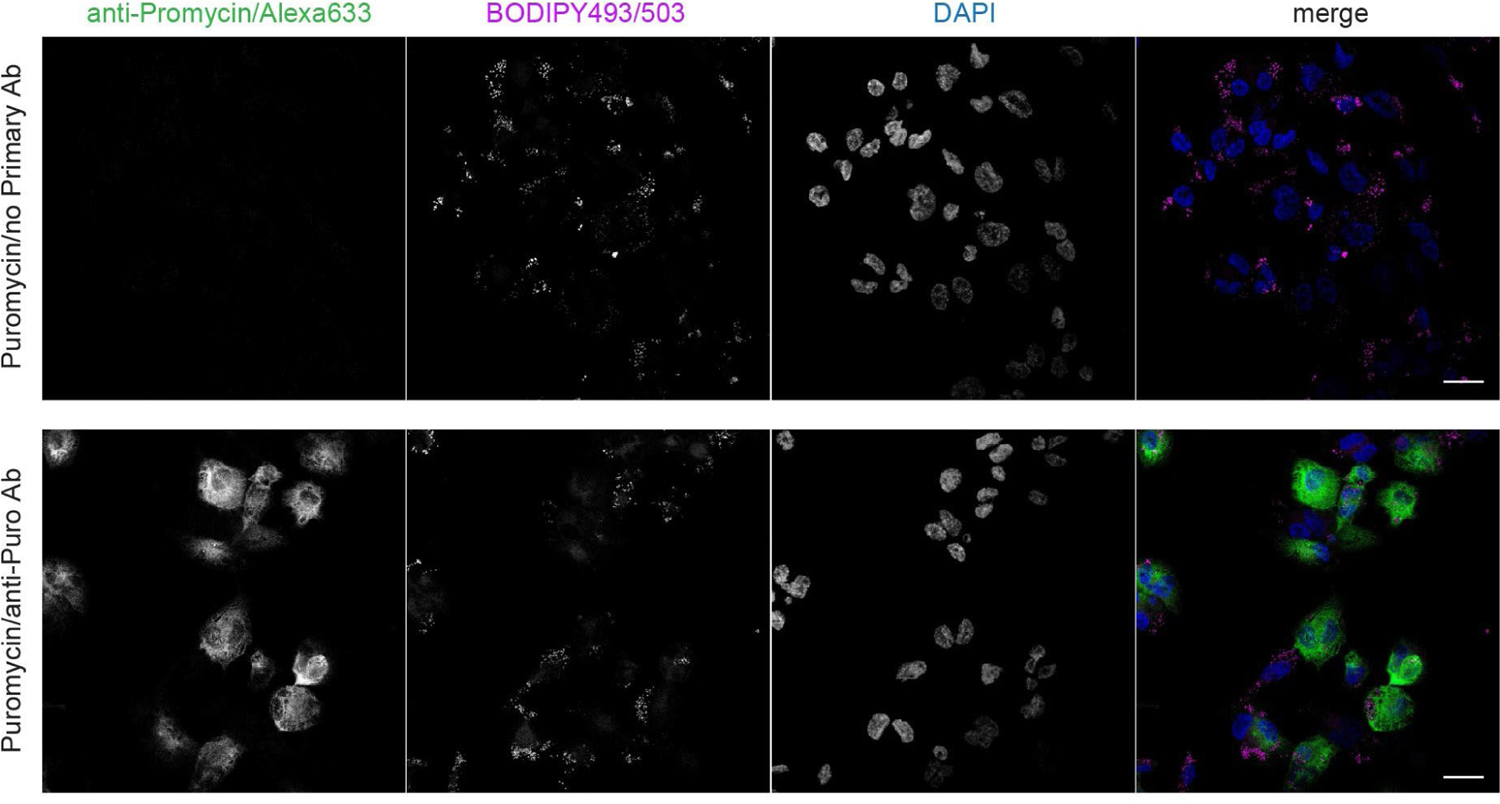
Immunofluorescence of puromycin treated THP1 macrophages with and without anti-puromycin primary antibody. Scale bar=10 μ

**Supplementary Figure 3.**
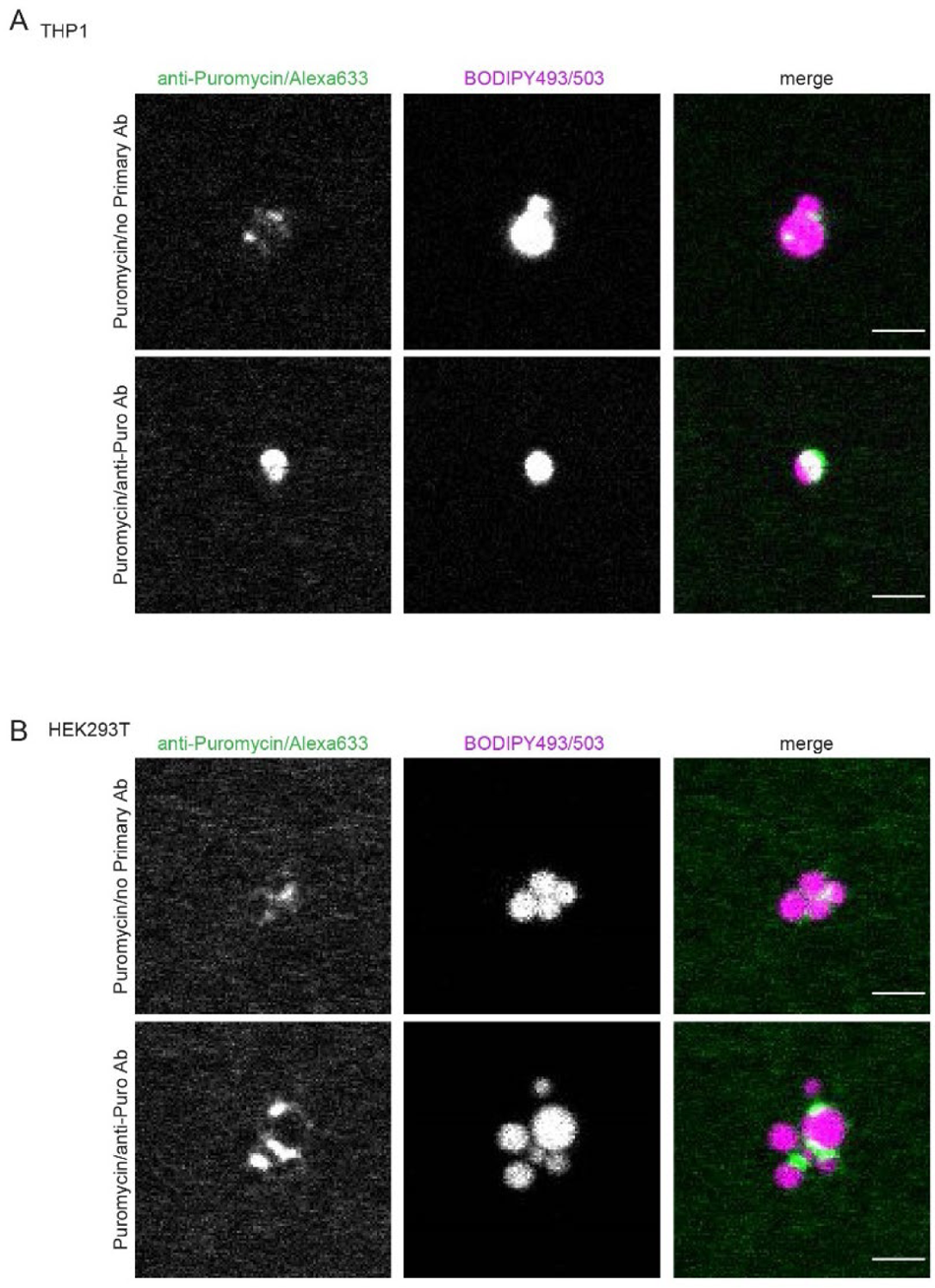
Immunofluorescence of LDs isolated from puromycin treated or untreated THP1 macrophages (A) or HEK293T cells (B) stained with Alexa633-tagged anti-puromycin antibody. Scale bar=2 μ

**Supplementary data Table 1.**
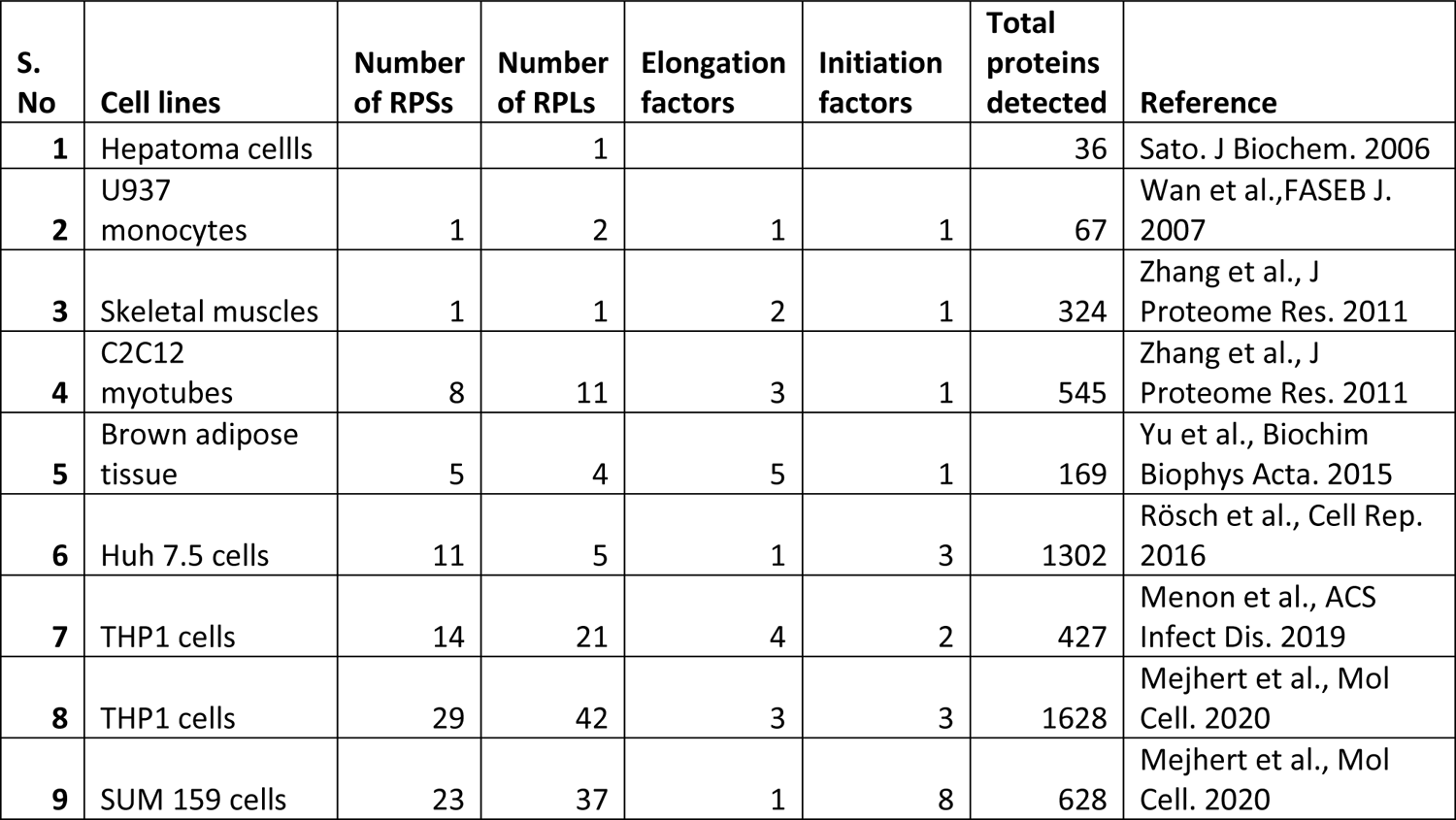

